# Age and IFN-β-induced changes in glial morphometry can be captured by in vivo diffusion-weighted magnetic resonance spectroscopy

**DOI:** 10.64898/2026.03.24.713975

**Authors:** Eva Periche-Tomas, Itamar Ronen, Jonathan Underwood, John Evans, Claire MacIver, Helena Leach, Francesca Branzoli, Mara Cercignani, Neil A. Harrison

## Abstract

**Introduction:** Neuroinflammation is increasingly implicated in age-related cognitive decline, neurodegeneration and neuropsychiatric disorders. During systemic inflammation, microglia are rapidly activated, simultaneously changing their shape and releasing cytokines that perturb neuronal function. This change in glial morphology alters their intracellular diffusion properties and provides a potentially measurable signature of their activation state. Diffusion-weighted magnetic resonance spectroscopy (dMRS) shows promise in detecting these changes. Here, we combined IFN-β challenge with dMRS to assess changes in metabolite diffusion in healthy young and older adults. We hypothesised that IFN-β would increase diffusion of choline-containing compounds (tCho) but not N-acetylaspartate + N-acetylaspartylglutamate (tNAA), age would be associated with an increase in tCho diffusion and concentration, lower tNAA concentration and increased effects of IFN-β.

**Methods:** We recruited 15 young (mean 25.2 ± 5.1 years, 6 male) and 15 older (mean 62.6 ± 4.1 years, 5 male) healthy volunteers, each tested twice, once after IFN-β and once after placebo. Physiological and behavioural responses were recorded hourly, and blood samples taken at baseline, 4 and 6.5 hours post-injection. dMRS occurred at ∼4.5 hours at 3T, using a semi-LASER sequence with four diffusion weightings (b = 0 and 3 × 3500 s/mm²), in 4.5 cm³ VOIs in the left thalamus and corona radiata. Apparent Diffusion Coefficients (ADCs) of tCho, tNAA and creatine+phosphocreatine (tCr) were calculated from averaged spectra using custom MATLAB software.

**Results:** IFN-β administration produced a significant increase in thalamic tCho diffusivity compared with placebo (t(28) = –2.15, p = 0.040), with no change in tNAA or tCr ADC, or tCho concentrations. IFN-β-related increases in tCho ADC positively correlated with increases in circulating IL-6 (R² = 0.14, p = 0.040). Age-related effects were also evident during the placebo condition, with older participants showing lower thalamic tNAA diffusivity (t(27) = 2.86, p = 0.008), lower tNAA/tCr in both grey and white matter (grey: t(27) = 2.49, p = 0.023; white: t(27) = 2.94, p = 0.007), and higher white-matter tCho/tCr (t(27) = –2.23, p = 0.034).

**Conclusion:** dMRS detected IFN-β–induced increases in thalamic tCho diffusivity corresponding with peripheral inflammation, supporting its sensitivity to acute inflammation-induced changes in glial morphology. Age-related differences in tNAA diffusion and concentrations further highlight metabolite-specific ageing effects.

**Highlights:**

- dMRS detects increased thalamic total choline diffusivity following IFN-β–induced inflammation.
- IFN-β–related changes in total choline diffusivity are associated with peripheral IL-6 responses.
- Ageing is linked to reduced NAA diffusion and higher white-matter tCho/tCr
- dMRS is sensitive to inflammation- and age-related neurochemical changes in vivo.

## INTRODUCTION

Glia are the most abundant and widely distributed non-neuronal cells in the central nervous system (CNS) and increasingly implicated in the aetiology of common mental illnesses, age-related cognitive decline and neurodegenerative disorders (Dar et al., 2024; Öngür et al., 1998; Rajkowska et al., 1999). As the brain’s resident immune cells, microglia constantly survey the parenchyma and rapidly respond to injury, infection, or systemic inflammation (Nimmerjahn et al., 2005). When reactive, they undergo coordinated morphological and functional changes, including retraction and thickening of processes and enlargement of their cell body, alongside increased expression of surface markers, release of pro-inflammatory cytokines, and enhanced phagocytic activity (Davalos et al., 2005; Kreutzberg, 1996). Like microglia, astrocytes, which play critical roles in the regulation of neuronal metabolism, synaptogenesis, ion homeostasis, and blood-brain barrier integrity, are also sensitive to systemic inflammation (Kucukdereli et al., 2011; Pellerin & Magistretti, 1994).

Like other cells, microglia and astrocytes’ physiological support roles are vulnerable to ageing, with changes in homeostatic function and phenotype commonly observed across the brain (Hanslik et al., 2021; Pan et al., 2020). For instance, during ageing, microglia show decreased cellular complexity and impaired injury response, together with increased mitochondrial activity and pro-inflammatory cytokine release. Astrocytes show similar changes in cellular complexity and mitochondrial activity, which is associated with impaired glutamate clearance (Hanslik et al., 2021). In rodents, age-dependent changes in microglial responses to intraperitoneal LPS have been reported from middle age (Keane et al., 2021; Nikodemova et al., 2016; Sierra et al., 2007), while non-interventional studies show that aged rodents display region-specific changes in microglial morphology, including a less ramified morphology with reduced process length and increased soma volume (Hefendehl et al., 2014; Tremblay et al., 2012). Astrocytes also undergo age-related morphological changes, with reduced size and short chubby processes, decreased astrocyte coupling through gap junction and variable changes in cell complexity (Bondi et al., 2021; Popov et al., 2021; Rodríguez et al., 2014).

These cellular changes are paralleled by shifts in brain chemistry measurable noninvasively through magnetic resonance spectroscopy (MRS). For example, ageing is associated with regional reductions in N-acetylaspartate (NAA), γ-aminobutyric acid (GABA), and the glutamate–glutamine complex (Glx). While NAA and GABA are largely neuronal, Glx reflects combined neuronal (glutamate) and glial (glutamine) metabolism via the glutamate–glutamine cycle (Rodríguez-Nieto et al., 2023; Hermans et al., 2018). Conversely, concentrations of glial-associated metabolites, including choline compounds (tCho) and myo-inositol (Ins), tend to increase with age (Rodríguez-Nieto et al., 2023). These patterns further underscore the complex and dynamic nature of glial and neuronal interactions across the lifespan.

Imaging glial responses in humans in-vivo remains challenging. To date, PET imaging using TSPO (18kDa translocator protein) tracers has been the predominant approach, based on the observation that TSPO expression increases with microglial activation. Several studies have demonstrated elevated TSPO binding following systemic LPS administration in humans and non-human primates (Sandiego et al., 2015; Hannestad et al., 2012). However, this method is limited by several considerations: it lacks cellular specificity (being expressed not only in microglia but also in astrocytes and endothelial cells) and is also sensitive to neuronal activity. In addition, a common TSPO polymorphism (rs6971) gives rise to high-, mixed-, and low-affinity binders, necessitating genotyping prior to PET imaging and complicating recruitment, quantification, and interpretation. This is particularly problematic in neuropsychiatric populations, where genotype distributions may themselves be biologically meaningful (Owen et al., 2012; (Notter et al., 2021; Owen et al., 2012). Moreover, recent evidence suggests that in humans, increases in TSPO binding predominantly reflect changes in microglial density rather than a shift to a reactive phenotype (Nutma et al., 2023). Compounding this, TSPO-PET is expensive, invasive, involves exposure to ionising radiation, and likely confounded by changes in peripheral binding during systemic inflammation, limiting its wider use (Schubert et al., 2021; Zhang et al., 2021). This has underscored a drive to find alternative MRI-based measures to directly or indirectly measure neuroinflammation.

Furthermore, fMRI and resting-state fMRI have been used to investigate effects of systemic inflammation on brain function and functional connectivity (Dipasquale et al., 2016; Kitzbichler et al., 2021), and quantitative magnetisation transfer (qMT) (Harrison et al., 2015) and diffusion-weighted MRI to assess effects on brain microstructure (Alexander et al., 2019). However, none of these techniques is directly sensitive to cell-type specific changes, limiting their ability to capture the distinct neuronal and glial responses that characterise neuroimmune responses during inflammation.

Diffusion-weighted magnetic resonance spectroscopy (dMRS) offers the potential to resolve this by exploiting the fact that some neurochemicals are differentially expressed in different cell types. For instance, total NAA (N-acetylaspartate + N-acetylaspartylglutamate) is neuronally restricted, while total choline, (tCho, including choline, phosphocholine, and glycerophosphocholine) and myo-inositol are predominantly expressed in glia. Total creatine (tCr, creatine + phosphocreatine) is present across all major cell types and can serve as a reference (Griffin et al., 2002; Urenjak et al., 1993; Gill et al., 1989). Because dMRS measures the apparent diffusion coefficient (ADC) of these metabolites, largely reflecting intracellular dynamics, it offers a potentially unique, non-invasive method for quantifying cell-specific microstructural changes in vivo (Palombo et al., 2018).

Consistent with this, dMRS has been shown to be sensitive to metabolite diffusion changes during inflammation. In mice, the Cuprizone mouse model of demyelination has shown elevated choline and myo-inositol ADCs in microglia and astrocytes, respectively, compared with control mice after 6 weeks of treatment (Genovese et al., 2021). Furthermore, these results correlated with histological measures of microglial and astrocytic alteration (Genovese et al., 2021). In humans, increased tCr and tCho diffusivity have been found in patients with amyotrophic lateral sclerosis (ALS) and systemic lupus erythematosus (SLE) and ischemic stroke (Ercan et al., 2016; Genovese et al., 2023; Reischauer et al., 2018; Zheng et al., 2012), together suggesting a pattern of glial activation.

Furthermore, increased choline diffusivity (but not NAA) has been observed in healthy controls after LPS peripheral administration (De Marco et al., 2022) and data from Multiple sclerosis (MS) patients have shown reduced thalamic NAA, which would be consistent with the neuronal damage and cell loss that are characteristic of the progression of autoimmune diseases (Bodini et al., 2018).

Here, we build on a recently validated model of mild systemic inflammation using subcutaneous administration of interferon-β (IFN-β) in healthy human participants (Periche-Tomas et al., 2026). This model induces a controlled inflammatory response characterised by elevated transient physiological, cellular immune, cytokine and behavioural responses, while avoiding the confounding cardiovascular effects often seen with higher-dose or intravenous challenges. This makes it an ecologically valid model for studying brain-specific immune responses, suitable for application across a broader population, including both younger and older adults.

Using dMRS, we assessed the effects of IFN-β on the diffusion properties of total NAA, choline, and creatine in the thalamus and corona radiata, representative regions of grey and white matter, respectively. These regions were selected based on their anatomical homogeneity, sensitivity to systemic immune activation, and high baseline TSPO expression in the case of the thalamus (Buttini et al., 1996; De Marco et al., 2022; Schubert et al., 2021).

We hypothesised that IFN-β would increase tCho ADC, consistent with glial reactivity, without significantly affecting NAA, which is primarily neuronal. We further predicted that these effects would correlate with peripheral inflammatory markers, particularly pro-inflammatory cytokines such as IL-6, and that age would modulate the IFN-β response, given known age-related alterations in glial function. By combining this novel experimental model of inflammation with a non-invasive and cell-type-sensitive imaging technique, our work offers new insight into glial dynamics, neuroimmune interactions, and the influence of age on these processes in the healthy brain.

## METHODS

### Participants

Thirty healthy adults were enrolled and stratified into two age groups: 15 younger adults (6 male (40%); mean age = 25.2 ± 5.1 years) and 15 older adults (6 male (40%); mean age = 65.6 ± 4.5 years). Participants were recruited from the Cardiff area. Eligibility criteria required participants to be in good physical and mental health and to be non-smokers, as verified by medical history, clinical assessment, vital signs, and laboratory screening. Laboratory tests included renal, hepatic, and thyroid function, as well as a full blood count.

In the younger group, most participants identified as White (n = 13), with the remaining two identifying as Asian; all participants in the older group identified as White. Participants were instructed to abstain from alcohol and strenuous physical activity for 24 hours prior to each study visit.

The study received ethical approval from the London–Camden & King’s Cross Research Ethics Committee (reference: 20/LO/0239), and all participants provided written informed consent prior to participation.

### Experimental design and inflammatory challenge

The present study forms part of a larger experimental medicine protocol designed to characterise physiological, behavioural, immune, and neurobiological responses to a mild inflammatory challenge in healthy younger and older adults, described in detail elsewhere (Periche-Tomas et al., 2026). The study employed a randomised, placebo-controlled, repeated-measures cross-over design, in which each participant completed two experimental sessions separated by 2–6 weeks (mean inter-session interval = 28.2 days). At one session, participants received 0.4 mL of reconstituted interferon-β (IFN-β; EXTAVIA®, 100 µg), and at the other session, they received 0.4 mL of 0.9% saline placebo, with treatment order counterbalanced across participants. IFN-β administration reliably induced a transient systemic inflammatory response, as previously reported.

### Systemic physiological, behavioural, and immune measures

As part of the broader protocol, systemic physiological, behavioural, and immune measures were collected at baseline and repeatedly following injection. Body temperature, heart rate, and systolic and diastolic blood pressure were recorded at baseline and at six post-injection time points (1, 2, 3, 4, 5.5, and 6.5 hours).

Subjective ratings of mood, fatigue, and sickness were collected using the Profile of Mood States (POMS; (McNair et al., 1971), a fatigue visual analogue scale (fVAS; (Gift, 1989), and the Karolinska Sickness Questionnaire (SicknessQ; (Andreasson et al., 2018). These were administered at baseline and at five subsequent time points. Blood draws for full blood count, and cytokine analysis (IFN-β, IL-6, TNF-α and IL-10) were taken at baseline, 4 hours, and 6.5 hours post-injection. The present report focuses on dMRS outcomes and their associations with behavioural and systemic inflammatory markers. Physiological, behavioural, and immune measures were included solely to permit exploratory correlation analyses with dMRS outcomes; full methodological details and primary results for these measures are reported elsewhere (Periche-Tomas et al., 2026)

### Neuroimaging markers

Neuroimaging data were acquired between 4.5 and 5 hours after each injection. This timing was informed by earlier human studies of Type I interferons (primarily IFN-α), which show that behavioural and inflammatory responses, such as changes in mood, fatigue, and motivation, emerge with approximately 4 hours of administration, indicating a robust peripheral and central inflammatory response (Davies et al., 2020; Dowell et al., 2016). While peak responses may occur later, this time window was selected to capture a robust early-phase response within a timeframe that was practical for study procedures and participant testing.

### MRI and MRS acquisition and analysis

For this study, MRI and MRS data were obtained at 3T (Siemens Magnetom Prisma, Siemens Healthineers, Erlanger, Germany). After 3-plane localizer, a T1-weighted MPRAGE sequence (magnetization prepared rapid acquisition gradient echo) was acquired in sagittal orientation and reconstructed in 3 orthogonal planes (TR = 2100 ms, TE = 3.24 ms, TI = 850 ms, flip angle = 8 degrees, FoV 256 x 256 mm^2^). These scans were used to position two 4.5 cm^3^ dMRS volumes of interest (VOIs) on the left thalamus and left corona radiata (20 x 15 x 15 mm^3^ voxel size). These two regions were selected based on (i) the need to capture signal from homogeneous grey and white matter brain areas (ii) that the left hemisphere is more commonly implicated in inflammation-induced cognitive disturbance (Haroon et al., 2014; Harrison, 2017).

The dMRS sequence used was a bipolar sequence based on a semi-Localization by Adiabatic SElective Refocusing sequence (semi-LASER) (Genovese et al., 2021) with TE = 100 ms, TR = 3 s, spectral width = 2500 Hz, number of complex points = 1024. Four diffusion weighting conditions were applied: one at b = 0 mm^2^/s and three at b = 3500 mm^2^ /s and diffusion gradients applied in three orthogonal directions [1, 1, -0.5], [1, -0.5, 1], [-0.5, 1, 1] in the VOI coordinate system.

For each condition, the number of signal averages (NSA) was 32 and a short scan without water suppression (NSA = 4) was performed for eddy current correction. B_0_ homogeneity was achieved by employing a rapid automated shimming method that utilised echo-planar signals sequences and incorporated mapping along projections, referred to as FAST (EST) MAP (Gruetter & Tkáč, 2000).

Spectra data were transferred and analysed with customised software to pre-process dMRS data implemented in Matlab R2021b (Mathworks, Natick MA, USA) previously described and validated for dMRS analyses (De Marco et al., 2022). Data pre-processing consisted of three main steps: (i) generation of eddy current correction (ECC) files based on water data acquisition and (ii) application of ECC, phase and frequency corrections to individual transients prior to averaging, yielding output spectra files ready for analysis. For the last step, (iii) the analysis and quantification of spectral data, linear prediction singular value decomposition (LPSVD) was performed. The peak amplitude area estimates for the three orthogonal directions (at high b values) were averaged and used to compute the ADC for the three metabolites of interest (tCho, tNAA and tCr). Figure 1 illustrates spectral data acquired in the grey and white matter regions of the same participant under the two conditions (placebo and IFN-β).

**Figure 1.**
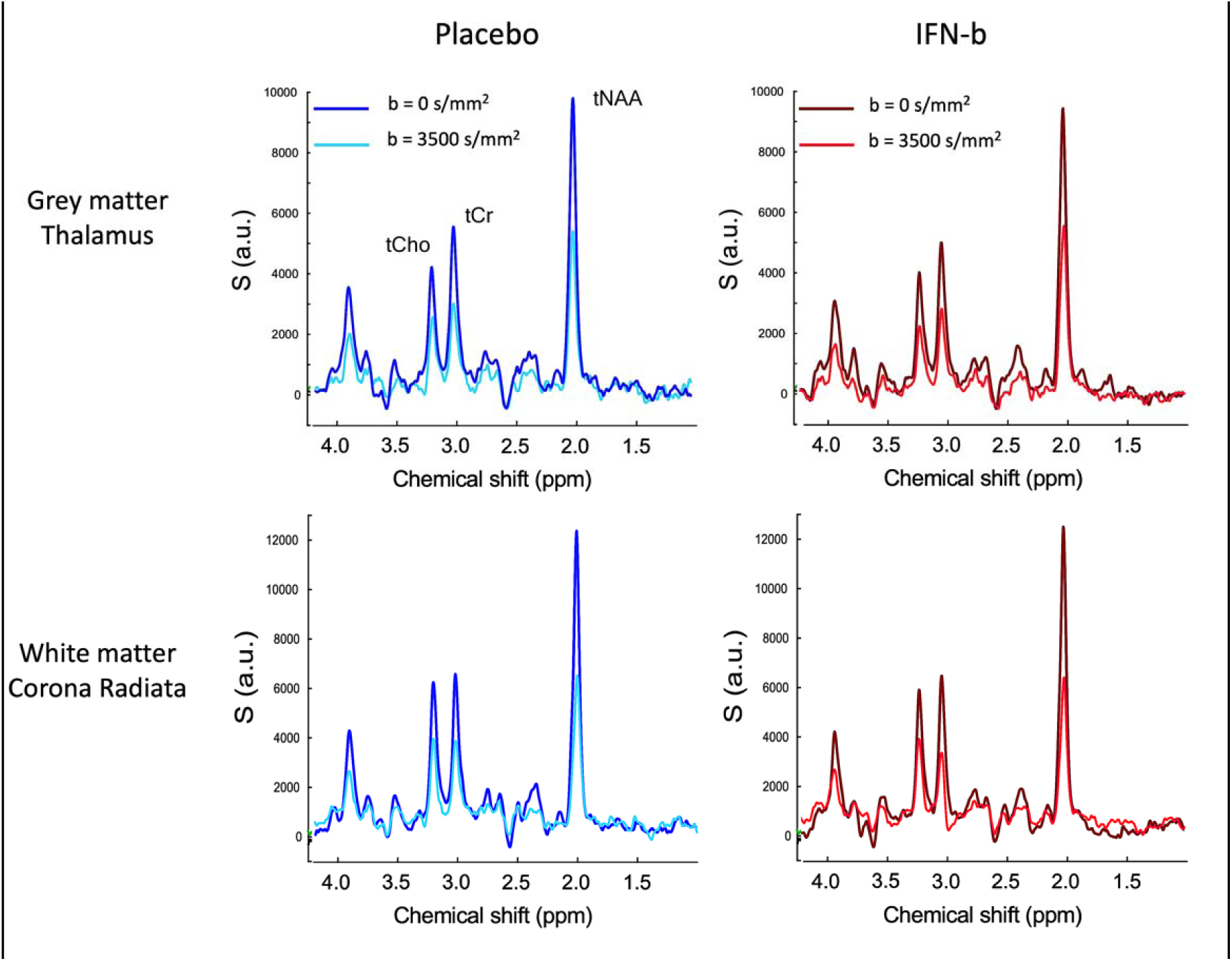
Example of MR spectra. Data were acquired at b = 0 mm^2^/s and b = 3500 mm^2^/s in the grey and white matter of one participant for the Placebo (blue) and IFN-β (red) conditions.

Amplitude values at b = 0 mm^2^ /s were used to estimate relative tCho and tNAA (concentrations, calculated as the ratio between their peak area and the tCr peak area). dMRS data were analysed completely blind to condition. Data from one participant was excluded due to a Cramer–Rao lower bound (CRLB) exceeding 10%, indicating low reliability of the ADC estimates. Consequently, the results section will present findings from 29 participants.

### Statistical Analysis

Paired-samples t-tests were used to compare apparent diffusion coefficient (ADC) values of total choline (tCho), total N-acetylaspartate (tNAA), and total creatine (tCr), as well as relative metabolite concentrations (tCho/tCr and tNAA/tCr), between the IFN-β and placebo conditions. Mixed factorial ANOVAs were conducted to assess condition × age group interactions on these measures. Age-related differences under placebo were evaluated using independent-samples t-tests. Sample size was determined a priori based on preliminary dMRS data demonstrating changes in choline diffusivity following inflammatory challenge (De Marco et al., 2022), which indicated that a sample of approximately 15 participants per group would provide adequate power to detect condition effects and condition x age interactions.

Pearson correlation coefficients were calculated to examine associations between IFN-β–induced changes in dMRS measures and changes in physiological indices, behavioural measures, and circulating cytokines. All statistical analyses were conducted using IBM SPSS Statistics version 27.

## RESULTS

### dMRS effects of IFN-**β** and age

#### Metabolite diffusion (Apparent Diffusion Coefficients)

Paired-samples t-tests revealed a significant increase in total choline (tCho) apparent diffusion coefficient (ADC) in the grey matter region of the thalamus following IFN-β administration compared with placebo (t(28) = 2.15, p = 0.040). Mean thalamic tCho ADC values were 1.43 × 10⁻L mm²/s (SD = 3.96 × 10⁻L) under placebo and 1.61 × 10⁻L mm²/s (SD = 4.52 × 10⁻L) following IFN-β administration.

No significant between-condition differences were observed for tCho ADC in the white matter region (corona radiata). Likewise, ADCs of tNAA and tCr did not differ significantly between IFN-β and placebo conditions in either grey or white matter regions (p>0.05; see Table 1). These data are summarised in Table 1. Mixed factorial ANOVAs did not reveal significant condition × age group interactions or main effects of age for any metabolite ADCs.

**Table 1.** Effects of IFN-β on metabolites ADC. Data represent mean ± SEM

| Mean<br>ADC mm<sup>2</sup>/s | Volume of<br>Interest | Condition |  |  |  |
| --- | --- | --- | --- | --- | --- |
|  |  | Saline | IFN-β | Δ% | p-values<br>(Saline vs. IFN- β) |
| tCho | Thalamus | 1.432e-04<br>(7.362e-06) | 1.614e-04<br>(8.393e-06) | 19.53<br>(7.85) | 0.04 |
|  | White Matter | 1.486e-04<br>(4.208e-06) | 1.492e-04<br>(4.113e-06) | 2.19<br>(3.68) | 0.894 |
| tNAA | Thalamus | 1.821e-04<br>(6.231e-06) | 1.910e-04<br>(7.563e-06) | 5.71<br>(3.2) | 0.143 |
|  | White Matter | 1.754e-04<br>(4.344e-06) | 1.791e-04<br>(4.129e-06) | 2.94<br>(2.39) | 0.366 |
| tCr | Thalamus | 1.644e-04<br>(6.244e-06) | 1.799e-04<br>(8.145e-06) | 13.61<br>(6.39) | 0.127 |
|  | White Matter | 1.754e-04<br>(4.344e-06) | 1.859e-04<br>(5.492e-06) | 7.124<br>(3.37) | 0.069 |

Distributions of ADC values under placebo and IFN-β conditions for each region are shown in Figure 2a–b. Between-condition differences in metabolite ADCs (IFN-β minus placebo) for the left thalamus are illustrated in Figure 3, and mean ADC values for all metabolites are reported in Table 1. We found no significant correlation between changes in body temperature induced by IFN-β and ADCs for any of the metabolites (all p> 0.1), indicating that metabolite diffusivity measures were not measurably related to systemic temperature variation.

**Figure 2.**
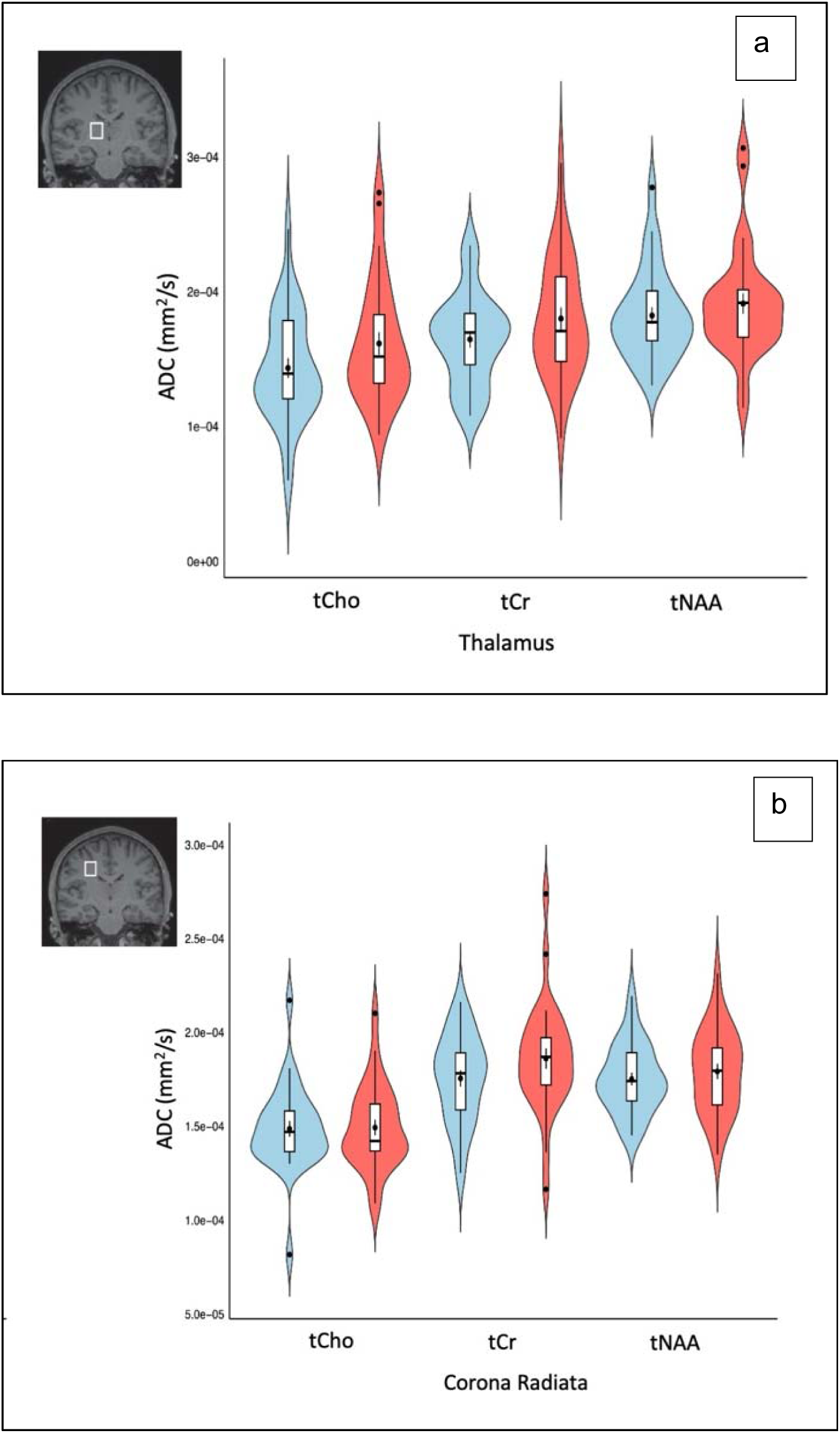
Grey and white matter metabolite ADC distribution. Violin plot illustrating data distribution, mean, median and interquartile ranges for the ADCs of (a) the grey matter (thalamus) and (b) white matter area (corona radiata) of the brain. Blue denotes placebo, red denotes interferon.

**Figure 3.**
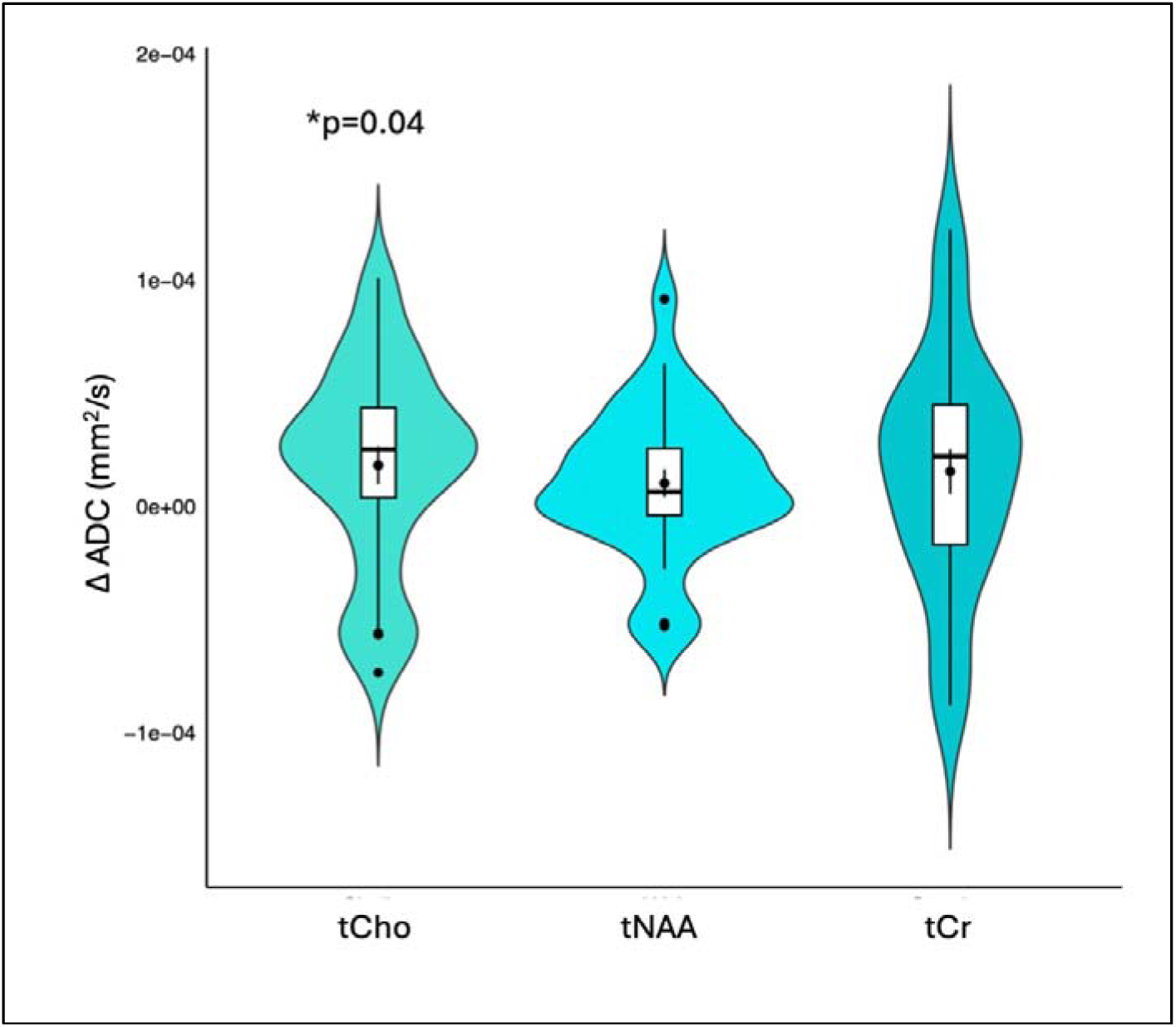
Grey matter metabolite ADC response to IFN-β. Differences between IFN-β and placebo sessions in the grey matter area of the brain. Mean, median and interquartile ranges are reported. P-value shows between-condition differences (IFN-β minus placebo) for choline ADC.

#### Metabolite concentrations

No significant between-condition differences were observed in relative tCho concentration (tCho/tCr) in either the thalamus or corona radiata, nor in thalamic relative tNAA concentration (tNAA/tCr). Mixed factorial ANOVAs did not reveal significant age effects for these measures.

In contrast, relative tNAA concentration in the white matter region (corona radiata) was significantly reduced following IFN-β administration compared with placebo (t(28) = −2.50, p = 0.016). Mean tNAA/tCr values were 2.15 (SD = 0.19) under placebo and 2.06 (SD = 0.24) following IFN-β administration (Figure 4a). Mixed factorial ANOVA revealed a significant condition × age group interaction (F(1,27) = 5.18, p = 0.031), as well as a significant main effect of age (F(1,27) = 21.52, p < 0.001), indicating lower relative tNAA concentrations in older participants . This effect appeared to be moderated by age and was further supported by an independent-samples t-test, which showed significantly lower relative tNAA concentrations in older compared with younger participants (t(27) = 2.95, p = 0.007; mean difference = 0.19, SE = 0.06) (Figure 4b).

**Figure 4.**
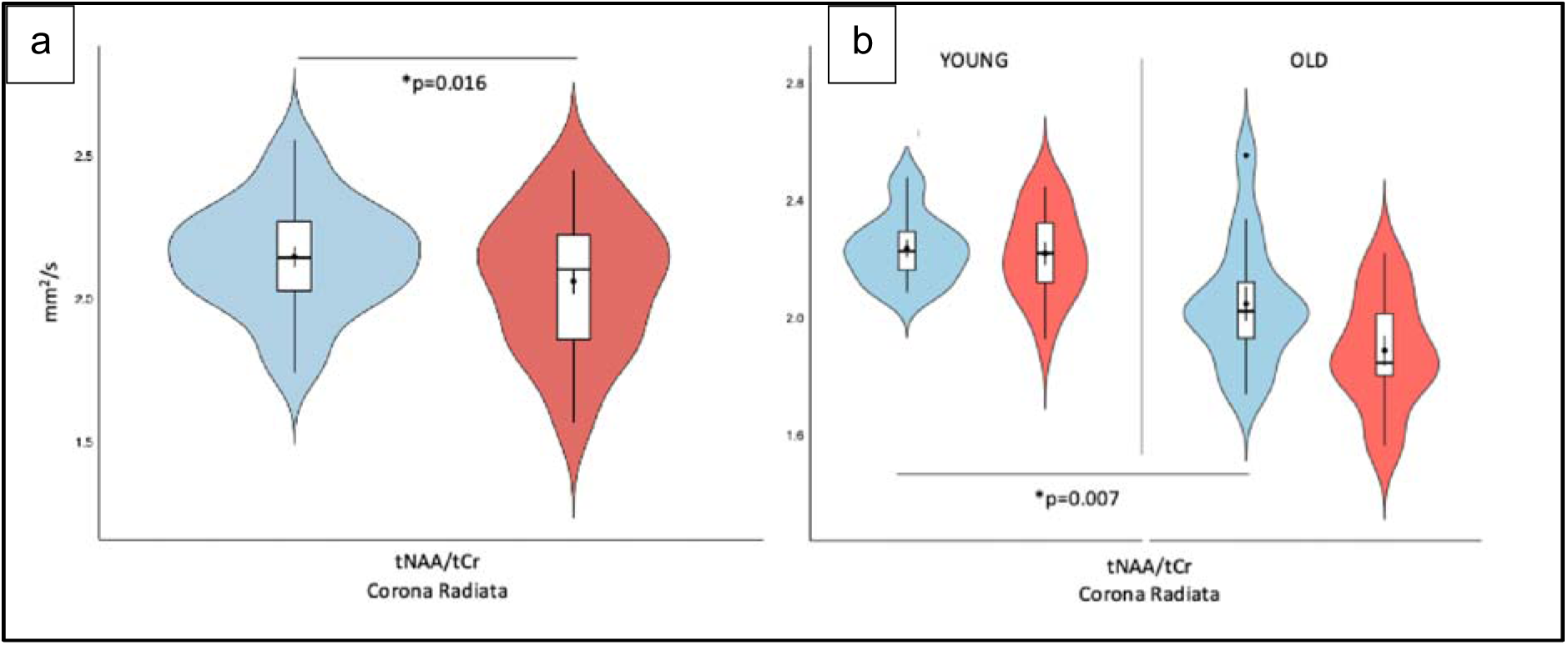
tNAA relative concentration across conditions and age groups. (a) Distribution of tNAA relative concentration ([tNAA/Cr]) for the IFN-β and placebo conditions in white matter. Significant values reflect main paired t-test results. (b) Distribution of tNAA relative concentration ([tNAA/Cr]) split by age group. Significant values reflect independent-samples t-test results at baseline. Blue denotes placebo; red denotes IFN-β.

#### Age-related differences in dMRS measures under placebo

To examine age-related differences independent of inflammatory challenge, dMRS measures were compared between younger and older adults under placebo conditions. Older participants showed significantly lower thalamic tNAA ADC compared with younger participants (t(27) = 2.86, p = 0.008; Figure 5a).

**Figure 5.** Age-associated differences in ADC and relative concentrations between young and old groups for the placebo condition. (a) tNAA ADC, (b) tNAA relative concentration in the Corona Radiata, (c) tNAA relative concentration in the Thalamus, (d) tCho relative concentration in the Corona Radiata. Significant values show reflect independent-samples t-test results. The young group is shown in blue, old group in green.

Relative tNAA concentration (tNAA/tCr) was also significantly lower in older adults in both white matter (t(27) = 2.94, p = 0.007; Figure 5b) and grey matter regions (t(27) = 2.42, p = 0.023; Figure 5c). In contrast, relative tCho concentration (tCho/tCr) in white matter was significantly higher in older participants compared their younger counterparts (t(27) = −2.23, p = 0.034; Figure 5d).

#### Associations between dMRS measures and behavioural and systemic markers

Potential associations between IL-6 levels and changes in tCho were assessed across the peripheral cytokines. ΔIL-6 was defined as the between-condition difference in within-session change (i.e., [6.5 hours − baseline] for IFN-β minus placebo). This measure was significantly associated with ΔtCho ADC (IFN-β minus placebo), such that greater IFN-β–related increases in IL-6 were correlated with increases in thalamic tCho ADC (R² = 0.14, p = 0.04; Figure 6).

**Figure 6.**
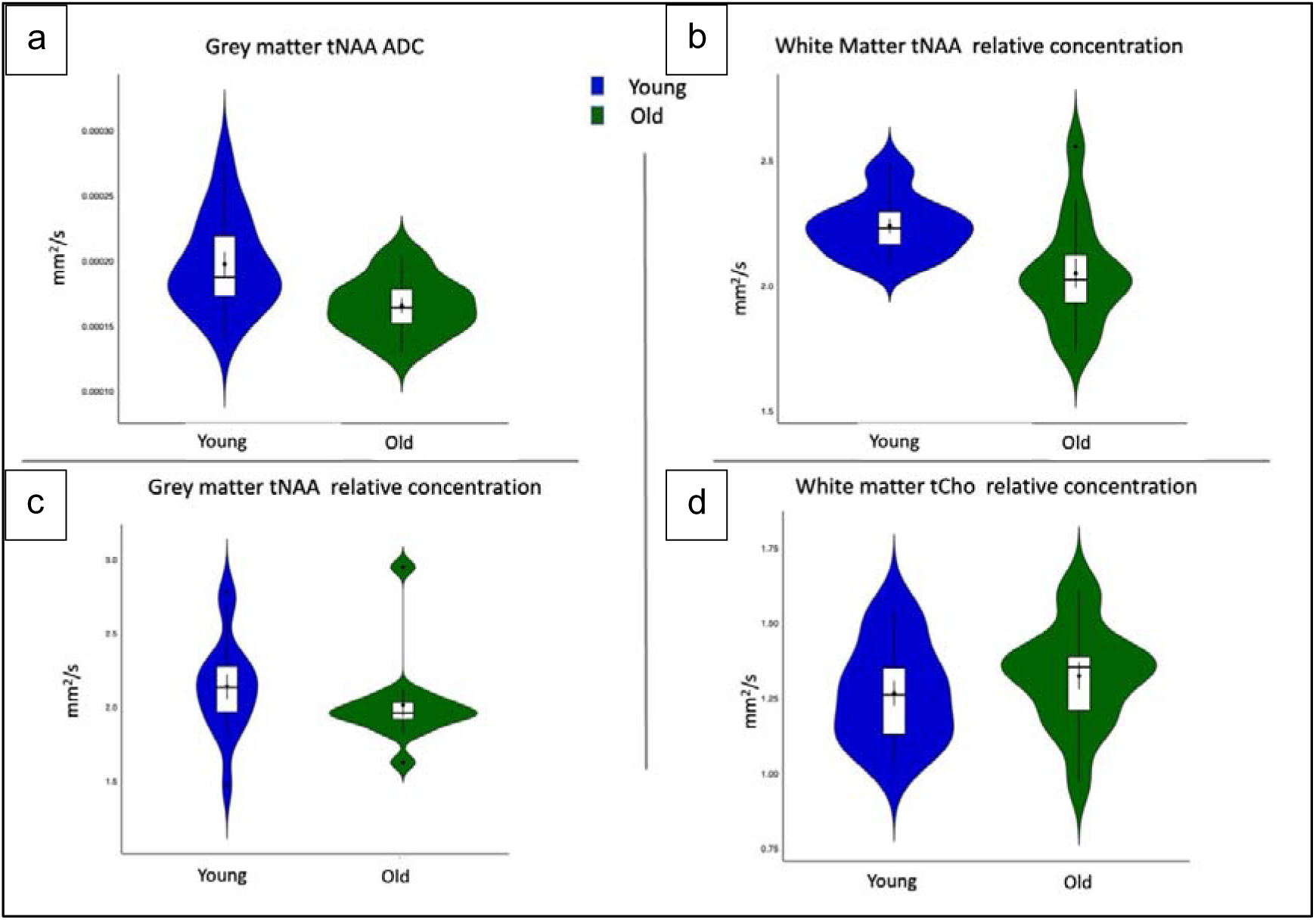
Correlation between tCho ADC and IL-6. Correlation between tCho ADC change between sessions and the difference between IL-6 plasma concentration levels (6.5 h post-injection minus baseline) in the two sessions. The inset shows IL-6 changes throughout the 2 sessions (blue denotes placebo, red interferon).

**Figure 7.**
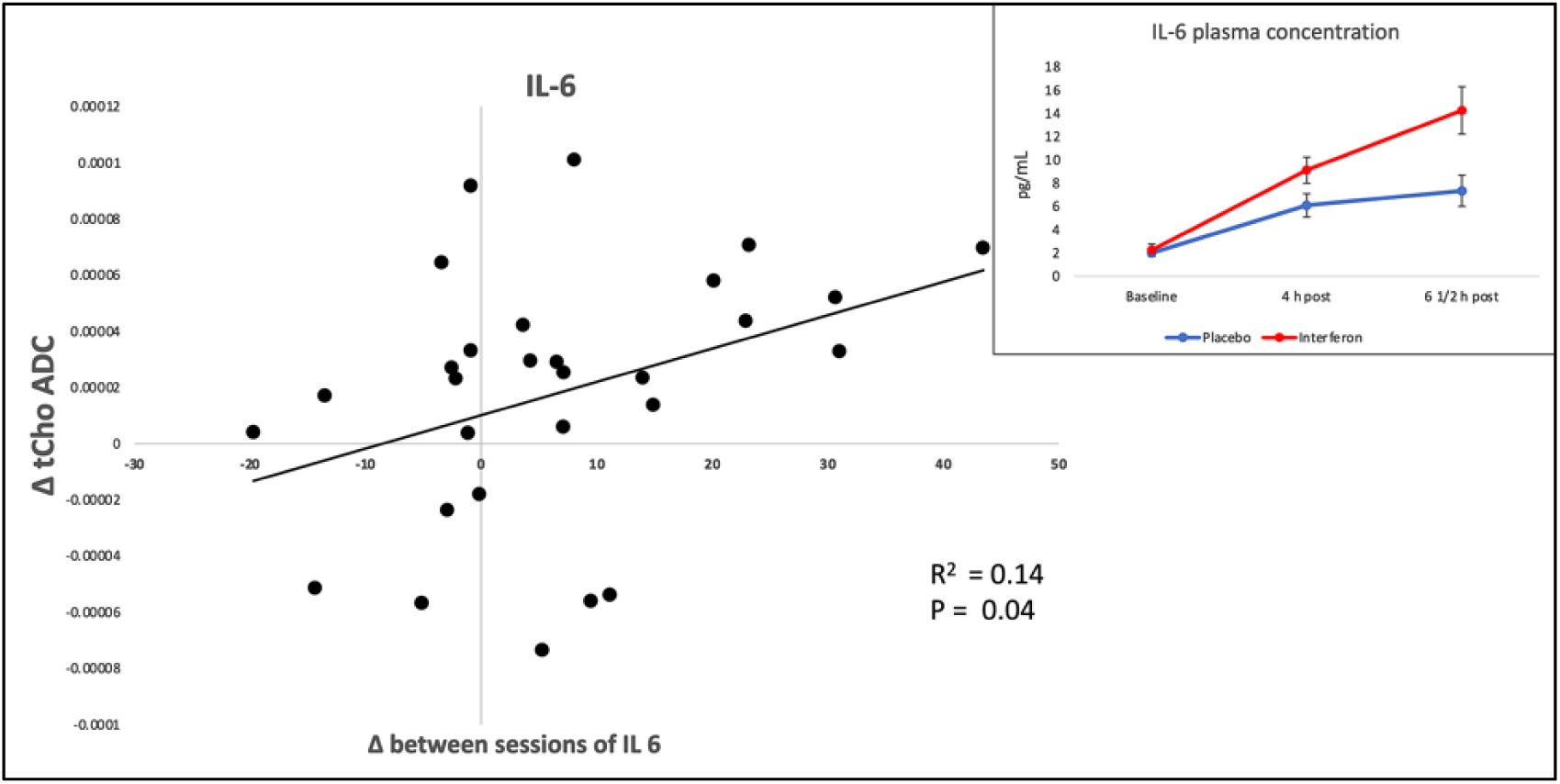
Correlation between tCho ADC and IL-6.

Exploratory analyses examining associations between IFN-β–induced changes in dMRS measures and physiological indices, behavioural measures (mood, fatigue, and sickness ratings), and cellular immune responses did not reveal any significant correlations (all p > 0.1).

## DISCUSSION

Here, we used dMRS in 15 young and 15 older adults to investigate the effects of experimentally induced inflammation on brain metabolite diffusion and concentration in healthy adult humans. We show that IFN-β administration is associated with changes in thalamic choline-compounds diffusivity, consistent with inflammation-related alterations in glial microstructure that can be detected *in vivo* using dMRS. We further demonstrate age-related differences in neuronal metabolite diffusion and concentration, indicating that ageing shapes the intracellular neurochemical environment and may influence vulnerability to inflammatory challenge. Finally, exploratory analyses suggest that individual variability in peripheral inflammatory responses (indexed by IL-6) relates to the magnitude of central choline diffusion changes. Together, these findings support dMRS as a sensitive tool for probing glial and inflammation-related microstructural and metabolic alterations in the human brain and highlight age-dependent neurochemical profiles.

The primary finding is consistent with experimental evidence showing that systemic inflammation induces cytomorphological changes in glial cells that alter intracellular metabolite diffusion (Verkhratsky et al., 2014). Prior dMRS studies in animal models have demonstrated increased choline and myo-inositol diffusivity associated with microglial and astrocytic alterations, respectively (Genovese et al., 2021), supporting the interpretation of tCho ADC as a marker of glial microstructural change.

Extending this work to humans, increased choline diffusivity has previously been reported in inflammatory neurological conditions such as systemic lupus erythematosus and amyotrophic lateral sclerosis (Ercan et al., 2016; Reischauer et al., 2018). In the context of experimental immune challenges, a preliminary dMRS study in healthy participants showed increased thalamic choline diffusivity following LPS administration (De Marco et al., 2022). Consistent with this literature, we observed increased thalamic choline diffusivity following IFN-β administration, although the magnitude of the effect was smaller than that reported after LPS, which is expected given the milder and more transient inflammatory response induced by IFN-β (Periche-Tomas et al., 2026).

Notably, both studies identified the thalamus as a sensitive region, with no detectable effects in white matter tracts such as the corona radiata, suggesting either greater resilience of myelinated pathways to short-lived inflammatory challenges or current methodological limits in detecting subtle white matter microstructural changes with dMRS.

The observed increase in choline diffusivity following IFN-β administration further supports the notion that inflammation-related changes in intracellular metabolite diffusion reflect cell-type–specific microstructural alterations that can be detected with dMRS, extending previous findings to a milder and more ecologically valid model of systemic inflammation. Recent evidence suggests that, in humans, increases in TSPO signal may reflect changes in microglial density rather than shifts in activation state per se (Nutma et al., 2023), highlighting ongoing challenges in interpreting TSPO PET findings. In this context, dMRS may offer complementary information by providing sensitivity to intracellular microstructural changes associated with inflammatory processes and could therefore contribute alongside established PET approaches to a more comprehensive in vivo characterisation of neuroinflammation.

Although both microglia and astrocytes undergo metabolic and morphological changes during neuroinflammation (Escartin et al., 2019; Heneka et al., 2014), it is not possible to definitively determine which glial cell type underlies the observed changes in choline diffusion. However, evidence from dMRS studies in animal models suggests a degree of specificity, with choline diffusivity more closely associated with microglial alterations, whereas myo-inositol diffusivity appears more sensitive to astrocytic hypertrophy (Genovese et al., 2021; Ligneul et al., 2019). In the present study, myo-inositol diffusion could not be reliably quantified due to the longer echo time required for the acquisition protocol, which limits the detection of metabolites with shorter T2 relaxation times.

Experimental models also indicate a temporally structured neuroimmune response, in which microglial activation often precedes astrocytic reactivity (Garcia-Hernandez et al., 2020; Liddelow et al., 2017).

Taken together, these findings suggest that the most parsimonious interpretation of the present results is that the observed changes in thalamic choline diffusion primarily reflect microglial rather than astrocytic activation following IFN-β administration.

In contrast to choline, we did not observe significant changes in tNAA diffusivity following IFN-β administration. As tNAA is considered a marker of neuronal integrity, altered tNAA diffusion has been associated with neuronal damage and cell loss in conditions such as multiple sclerosis and cerebral ischemia (Bodini et al., 2018; Zheng et al., 2012). The absence of tNAA diffusivity changes in the present study, consistent with findings from other experimental inflammation models (De Marco et al., 2022) suggests that the mild inflammatory challenge induced by IFN-β does not produce measurable alterations in neuronal morphology.

Pronounced age-related effects were observed under placebo conditions: older adults showed reduced thalamic tNAA diffusivity and lower relative tNAA concentrations in both grey and white matter, alongside higher relative choline concentration in white matter. Notably, the reduction in white-matter relative tNAA concentration observed following IFN-β administration appeared to be moderated by age, with older participants showing lower baseline relative tNAA concentrations, suggesting that pre-existing age-related differences in neuronal metabolites may contribute to the observed difference between IFN-β and placebo conditions. These findings align with histological and neuroimaging evidence of age-associated reductions in neuronal density and dendritic complexity (Bishop et al., 2010; Morrison & Hof, 2003; Raz et al., 2004), which would be expected to restrict intracellular space available for metabolite diffusion and lower neuronal markers such as tNAA. Complementary work has reported region-specific increases in density and glial markers with age, together with elevated glial-associated metabolites including choline and myo-inositol (Deelchand et al., 2020; Martínez-Pinilla et al., 2016; Rodríguez et al., 2014; Rodríguez-Nieto et al., 2023). Taken together, the co-occurrence of lower tNAA diffusivity/concentration and higher white-matter choline in older adults supports the interpretation that dMRS is sensitive to age-related shifts in the neuronal–glial balance of the intracellular environment in vivo.

Contrary to our hypothesis, we did not observe significant condition-by-age interactions in choline diffusivity or other metabolite measures. This may reflect limited statistical power to detect subtle interaction effects, the relatively healthy status of the older cohort, or the mild and transient nature of the IFN-β inflammatory challenge. It is also possible that age-related differences in glial responsiveness emerge more clearly under conditions of heightened or chronic inflammation, or at different post-challenge time points not captured in the present study (Keane et al., 2021; Nikodemova et al., 2016; Şimşek et al., 2025). Notably, neuroimaging was conducted during an early post-injection window (4.5–5 hours), selected to capture a robust initial inflammatory response based on prior work, while remaining feasible within the constraints of the experimental protocol; age-related effects may therefore be more apparent at later phases of the response.

Exploratory analyses revealed a positive association between IFN-β–induced changes in plasma IL-6 and increases in thalamic choline diffusivity, linking peripheral inflammatory signalling to central neurochemical alterations. The thalamus is a grey matter region with high microglial density and sensitivity to inflammatory stimuli (Buttini et al., 1996; Schubert et al., 2021) and individuals mounting a larger peripheral inflammatory response may therefore exhibit more pronounced central effects. No significant associations were observed between choline diffusivity and behavioural or physiological measures. This dissociation is consistent with the broader literature on inflammation-associated behavioural change, which implicates limbic and mesolimbic circuits—such as the subgenual cingulate cortex, amygdala, and ventral striatum—rather than the thalamus, in the emergence of mood and sickness-related symptoms following immune challenge (Capuron et al., 2012; Harrison, Brydon, Walker, Gray, Steptoe, & Critchley, 2009; Harrison, Brydon, Walker, Gray, Steptoe, Dolan, et al., 2009). Together, these findings suggest that thalamic choline diffusion changes observed following IFN-β administration may reflect early or upstream neuroimmune processes that are sensitive to peripheral inflammatory signalling but are not directly coupled to behavioural expression under conditions of mild or transient immune activation.

Several methodological considerations are relevant when interpreting the present findings and in evaluating the suitability of dMRS for studying inflammation-related brain changes in humans. Due to practical and time constraints, data acquisition was restricted to two volumes of interest (VOIs), selected based on prior literature using LPS as the most commonly employed experimental model of systemic inflammation. Notably, microglial responses to different pro-inflammatory stimuli vary substantially, with LPS inducing a more robust pro- and anti-inflammatory transcriptional response compared with interferons (Lively & Schlichter, 2018). In this context, the thalamus represents a particularly sensitive grey matter region, exhibiting high baseline TSPO expression (Schubert et al., 2021) and consistent responsiveness to systemic immune challenge (Buttini et al., 1996). Diffusivity changes observed in this region may therefore reflect relatively general inflammation-related glial microstructural alterations in grey matter, rather than region-specific substrates of behavioural change. Future studies incorporating a broader range of VOIs will be important for establishing the regional specificity of dMRS markers and their relationship to behavioural and clinical phenotypes.

A potential confound in studies of inflammation-induced diffusion changes is the effect of elevated body temperature on metabolite diffusivity. In the present study, IFN-β administration was associated with increases in body temperature; however, we did not observe changes in tNAA diffusivity in either VOI, nor significant changes in tCho diffusivity in white matter. Importantly, no associations were found between peak temperature changes and metabolite ADCs, suggesting that the observed effects are unlikely to be driven by non-specific thermal influences on diffusion.

Another limitation of the current design is that dMRS data were acquired at a single post-injection time point (approximately 4½–5 hours following administration), precluding direct assessment of the temporal evolution of neuroimmune responses. Previous work using LPS has shown that physiological responses peak within 2–3 hours, while TSPO PET studies demonstrate glial activation between 3–5 hours in humans and 4–6 hours in non-human primates (Hannestad et al., 2012; Sandiego et al., 2015).

In contrast, physiological, behavioural, and cellular immune responses in the present IFN-β model peaked later, at approximately 6.5 hours post-injection, suggesting a delayed inflammatory trajectory. These differences highlight the importance of stimulus-specific timing and indicate that dMRS may be well suited to capturing intermediate or evolving microstructural changes associated with milder inflammatory challenges. Future studies incorporating repeated imaging time points could clarify the temporal dynamics of glial activation and resolution and their relationship to acute and persistent symptoms.

From a technical perspective, dMRS acquisition and processing present several challenges, including sensitivity to motion and limitations in signal-to-noise ratio (SNR). Linear translational motion can introduce phase shifts that are correctable during post-processing, whereas non-linear motion sources, such as cardiac pulsation, can lead to signal drop-out and overestimation of diffusion coefficients. To mitigate these effects, the processing pipeline employed here incorporates stringent quality-control criteria based on variance in metabolite peak amplitudes, an approach that has been shown to improve the reliability of ADC estimates (Genovese et al., 2021). Although the inclusion of diffusion gradients necessitates longer echo times and constrains the number of signal averages, optimised dMRS protocols have demonstrated sufficient robustness and reproducibility to detect biologically meaningful diffusion changes in vivo at 3T (Wood et al., 2015). In the present study, diffusion weighting was limited to two b-values (b = 0 and 3,500 s/mm²), enabling estimation of ADC but not higher-order diffusion metrics. Future studies incorporating multiple and higher b-values could allow modelling of non-Gaussian diffusion behaviour, such as diffusion kurtosis, potentially providing additional sensitivity to microstructural complexity and intracellular heterogeneity.

Taken together, these considerations indicate that dMRS represents a feasible and sensitive approach for probing intracellular microstructural changes associated with mild systemic inflammation in humans. As methodological developments continue, dMRS may offer a non-invasive means of detecting inflammation-related neurochemical diffusion changes that may complement existing neuroimaging approaches and contribute to the in vivo characterisation of neuroimmune processes.

In summary, the present findings demonstrate that dMRS is sensitive to IFN-β–induced changes in thalamic choline diffusion, consistent with inflammation-related glial microstructural alterations that can be detected *in vivo* in humans using a mild experimental model of systemic inflammation. We additionally observed robust age-related differences in metabolite diffusion and concentration, indicating that dMRS can capture changes in the intracellular neurochemical environment associated with healthy ageing. No significant interactions between inflammatory conditions and age were detected.

Together, these findings support the feasibility of dMRS as a non-invasive approach for probing neuroimmune and age-related brain changes in humans. While the present results provide an important foundation, future work addressing methodological considerations such as sample size, regional coverage, and variability in inflammatory challenge may help to elucidate more subtle effects and clarify how ageing modulates neuroimmune responses.

## Data access

Data will be made available on request.

## Acknowledgements

This work was supported by the Hodge Centre for Translational Neuroscience JU was supported by the Medical Research Council (grant number MR/T023791/1) FB acknowledges support from the programs ’Institut des neurosciences translationnelle’ ANR-10-IAIHU-06 and ’Infrastructure d’avenir en Biologie Santé’ ANR-11-INBS-0006.

The authors would like to thank Dr. Edward J. Auerbach and Dr. Małgorzata Marjańska for providing us with the dMRS sequence for the Siemens platform and simulating the basis set for spectral fitting.

